# Flygenvectors: The spatial and temporal structure of neural activity across the fly brain

**DOI:** 10.1101/2021.09.25.461804

**Authors:** Evan S. Schaffer, Neeli Mishra, Matthew R. Whiteway, Wenze Li, Michelle B. Vancura, Jason Freedman, Kripa B. Patel, Venkatakaushik Voleti, Liam Paninski, Elizabeth M.C. Hillman, L.F. Abbott, Richard Axel

**Affiliations:** Mortimer B. Zuckerman Mind Brain Behavior Institute and Department of Neuroscience, Columbia University, New York, NY, 10027, USA; Department of Statistics and the Grossman Center for the Statistics of Mind, Columbia University, New York, NY, 10027, USA; Department of Biomedical Engineering, Columbia University, New York, NY 10027, USA; Department of Radiology, Columbia University, New York, NY 10027, USA; Department of Physiology and Cellular Biophysics, Columbia University, New York, NY 10032, USA; Department of Biochemistry and Molecular Biophysics, Columbia University, New York, NY 10032, USA; Howard Hughes Medical Institute, Columbia University, New York, NY 10027, USA

## Abstract

What are the spatial and temporal scales of brainwide neuronal activity, and how do activities at different scales interact? We used SCAPE microscopy to image a large fraction of the central brain of adult *Drosophila melanogaster* with high spatiotemporal resolution while flies engaged in a variety of behaviors, including running, grooming and flailing. This revealed neural representations of behavior on multiple spatial and temporal scales. The activity of most neurons across the brain correlated (or, in some cases, anticorrelated) with running and flailing over timescales that ranged from seconds to almost a minute. Grooming elicited a much weaker global response. Although these behaviors accounted for a large fraction of neural activity, residual activity not directly correlated with behavior was high dimensional. Many dimensions of the residual activity reflect the activity of small clusters of spatially organized neurons that may correspond to genetically defined cell types. These clusters participate in the global dynamics, indicating that neural activity reflects a combination of local and broadly distributed components. This suggests that microcircuits with highly specified functions are provided with knowledge of the larger context in which they operate, conferring a useful balance of specificity and flexibility.

## Introduction

What are the spatial and temporal scales of activity in the brain? In nematodes, flies, zebrafish and mice (1), the exogenous activation of small clusters of neurons can drive behavioral sequences, providing a causal link between the activity of small groups of cells and specific behaviors. In *Drosophila melanogaster*, the identification of local circuits that govern specific behaviors has suggested a view of the fly brain as a collection of highly specialized microcircuits. On the other hand, brainwide recording of neural activity in multiple organisms reveals global activity associated with behavior (2–16) and task-related variables (17–19). Moreover, the activity of non-motor circuits is modulated by behavior (20–25). How do these observations relate to the view of the brain as composed of functionally specialized microcircuits? What is the relationship between signals that are broadly distributed and those that are local?

The analysis of brainwide activity at both a global and local scale requires that we simultaneously observe the activity of neurons distributed throughout the brain at sufficient temporal resolution to reveal correlations between neurons. We used SCAPE microscopy (26, 27), a single-objective form of light-sheet microscopy that permits high-speed volumetric imaging, to record activity in a significant fraction of the neurons across a large and contiguous portion of the brain of behaving *Drosophila*. This provides a comprehensive picture of activity in the fly brain. The principal patterns of neural activity (flygenvectors) comprise multiple spatial and temporal scales. We observe that signals related to some but not all behaviors engage the majority of neurons throughout the brain. Moreover, although most neurons are correlated with current behavior, a significant fraction are correlated with behavioral dynamics on longer timescales.

Although a small number of behavior-related signals dominates global activity, neural activity is rich and high-dimensional. Most of these dimensions are sparse and spatially organized, consistent with each dimension corresponding to localized activity of specific cell types. Therefore, neural activity in the behaving fly reflects the coordination of broadly distributed and local dynamics.

## Results

### Brainwide functional imaging at single neuron resolution

We used SCAPE microscopy to examine activity across a large fraction of the central brain of behaving adult *Drosophila*. This enabled dual-color imaging of the dorsal third of the central brain in the behaving fly at more than 10 volumes per second (Fig. 1A-B, G). We imaged flies expressing the nuclear calcium reporter nuc-GCaMP6s and the static nuclear dsRed under control of the panneuronal driver NSyb-Gal4. Expression is poor in Kenyon cells and they were omitted from our analyses (Fig. S1). Electron microscopy of the fly brain reveals approximately 30,000 cells in the central brain (30). We imaged the dorsal third of the central brain, achieving single-cell resolution through the majority of this imaged volume. After accounting for heterogeneity in cell density in the brain, depth limits of cellular resolution due to scattering, and omission of Kenyon cells, we expected to resolve on the order of thousands of cells. We used the fluorescence of the static red channel to extract on average 1,631 ± 109 ROIs per animal. After refinement to exclude ROIs with large motion artifacts, we obtained 1,419 ± 78 stable, single-cell ROIs per animal (Methods).

**Figure 1.**
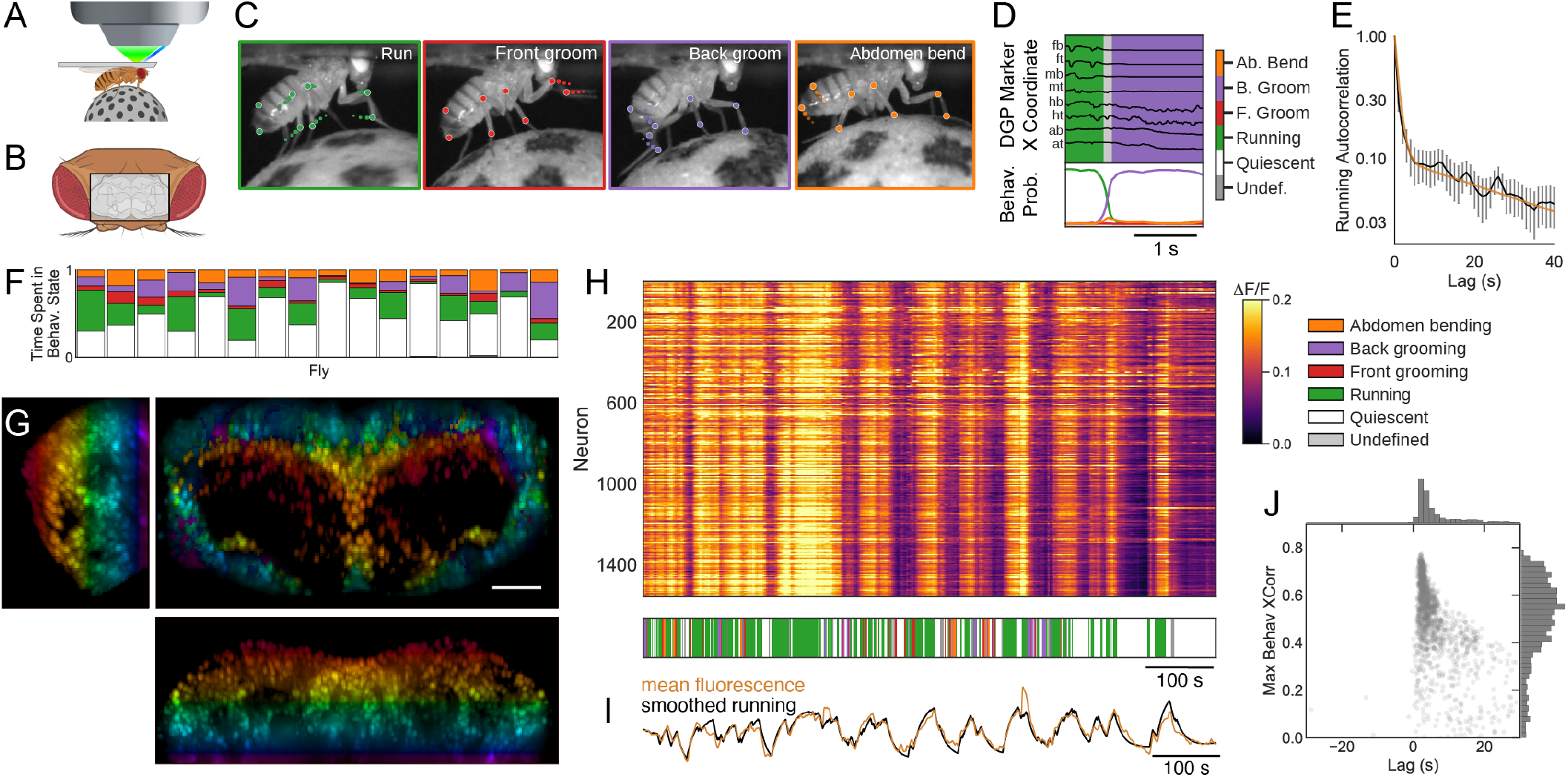
Brainwide neural activity correlates with behavior. (A) Illustration of SCAPE’s imaging geometry, in which an oblique light sheet (dark blue) sweeps across the fly brain and emitted light (green) is collected by the same objective lens. An image-splitter within the microscope permits dual-color imaging while the fly is head-fixed and running on a spherical treadmill. (B) The head of a fly viewed from a dorsal perspective (Top = posterior), with the approximate imaging window denoted by a black rectangle. (C) Points on the fly’s limbs and body are tracked with Deep Graph Pose (DGP) (28). Running, grooming and abdomen bending exhibit distinct patterns of limb dynamics, observed in trajectories of DGP points. (D) A semi-supervised sequence model (29) extracts a time series of discrete behavioral states from DGP points. Example trajectories of the 8 tracked points shown in black above, ordered from anterior to posterior (fb:front bottom, ft:front top, mb:middle bottom, mt:middle top, hb:hind bottom, ht:hind top, ab:abdomen bottom, at:abdomen top). Inferred probability of each behavioral state is shown below, showing a transition from running to back grooming. (E) The autocorrelation of running (black) is best fit by the sum of two exponentials, with time constants of 1s and 40s (gold). (F) Fraction of time spent in each behavioral state for each fly. Colors as in ‘D’. (G) Sample volume of raw imaging data in a brain with pan-neuronal expression of both nuclear-localized GCaMP6s and nuclear dsRed. Shown are maximum-intensity projections of the dsRed channel over the approximate dorsal/ventral (middle), anterior/posterior (bottom), and medial/lateral dimensions (left). Pseudocolor indicates depth in dorsal/ventral dimension. Scale bar in spatial map is 50 *μ*m. (H) *Top*, raster of ratiometric fluorescence for all neurons from one fly (Fly 1 in ‘F’). *Bottom*, behavioral state, color coded as in ‘D’. (I) Average ratiometric fluorescence from all neurons (gold) and running smoothed with an exponential filter (black, time constant = 6s) are highly correlated (r = 0.90). (J) Maximum cross-correlation with running for every cell from the same fly as in ‘H’, versus the corresponding lag. Each point is one cell.

### Broad-scale neural activity is highly correlated with behavior

We examined brainwide neural activity while flies behaved freely on a spherical treadmill (Methods). We identified the different behaviors exhibited by the fly by tracking points on the fly’s body with Deep Graph Pose (28). We used a semi-supervised approach (described in a companion manuscript (29)) to infer the behavioral states of running, front and back grooming, abdomen bending and quiescence (Fig. 1C-D, Methods). Different flies exhibited these behaviors with varying frequencies (Fig. 1F). We also imaged the fly without a spherical treadmill, where it primarily exhibited a flailing behavior. On the treadmill, flies performed bouts of running punctuated by either grooming or quiescence. Autocorrelation of the running state decayed on time scales of 1s and 40s (Fig. 1E), because running occurred in bouts that lasted a few seconds but the tendency to run persisted for considerably longer times. The other annotated behaviors exhibited only a single fast correlation time (Fig. S1). Long-timescale changes in the tendency to run suggest that an underlying state, such as arousal, fluctuated over the course of our experiments.

Strikingly, most of the imaged neurons throughout the brain show a pattern of activity that is correlated with running. This is in accord with previous studies demonstrating that most of the neuropil in the fly brain is active when the fly runs (6, 7) and demonstrates that these earlier neuropil recordings are not the consequence of a sparse ensemble of active neurons with extensive projections. Rather, running is represented by the vast majority of neurons in the fly brain (Fig. 1H). The mean activity across all the imaged neurons is highly correlated with running smoothed with an exponential filter with a decay time of 6s (r = 0.90, Fig. 1I). This correlation cannot be accounted for by motion artifacts (r = 0.02, Fig. S1). Cross-correlation of individual neurons with running is high, and the activity of most neurons follows running with a small lag (Fig. 1J).

### Distinct neural populations represent locomotion over different timescales

We fit a regression model to extract the components of neural activity correlated with the identified behaviors (running, front and back grooming, abdomen bending, and quiescence). The observation that the auto-correlation of running exhibited two decay times (Fig. 1E) suggested that different neurons might be correlated with behavior on different timescales. Therefore, we regressed each neuron’s activity against all behaviors filtered using a different fitted time constant (*τ*_*i*_) for each cell (*i*). We explored the causal relationship between behavior and neural activity by allowing for a cell-specific temporal shift (*ϕ*_*i*_) of neural activity relative to the annotated behaviors (Methods).

Regressing neurons across behaviors and filtering each neuron with its own time constant considerably increased correlations between the activity of individual neurons and the annotated behaviors (Fig. 2A). This model accounted for proportionally more variance in flies that spent more time running (*CC* = 0.73, Fig. 2B), as expected from the widespread representation of running (Fig. 1H). The majority of neurons are positively correlated with running, although a smaller population show strong negative correlation with running (Fig. 2C). Negatively correlated neurons are highly concentrated in the Pars Intercerebralis (PI) region (Fig. 2D-F), a heterogeneous population of peptidergic neurons involved in a wide range of functions (31).

**Figure 2.**
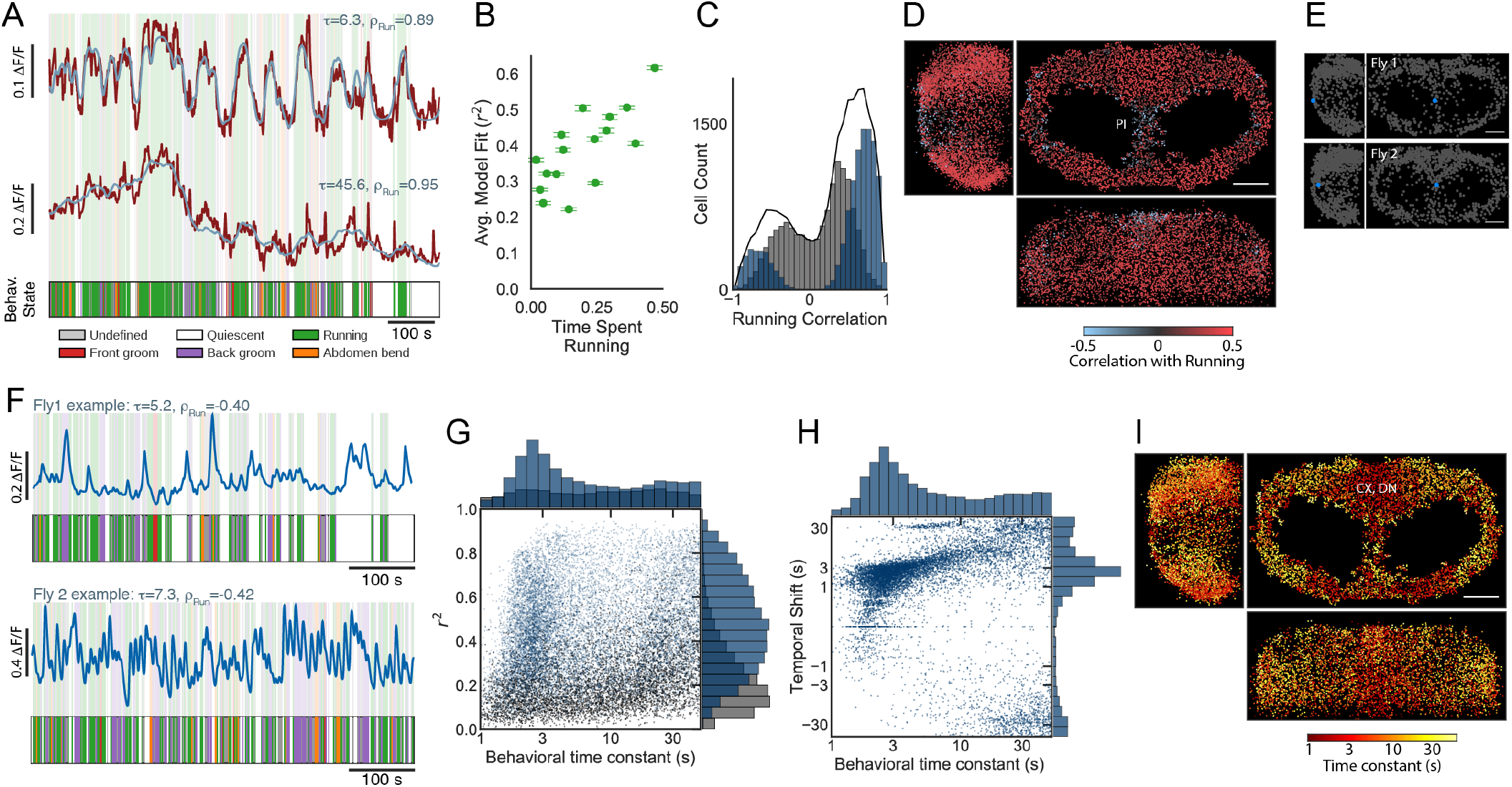
Brainwide correlates of running are multimodal and spatially organized. (A) Example traces from selected cells with small (top) and large (bottom) behavior time constants, with regression fit overlaid in blue and ethogram below. (B) Average model fit across cells for each fly versus fraction of time spent running (N = 16). Fly with highest time spent running shown in Figure 1H-J. (C) Correlation with running for all cells and all flies (N = 18), cells significantly active during behavior in blue, all other cells in gray, total in black. (D) Downsampled composite spatial map of running correlation for all flies, viewed in the sagittal (left), transverse (right), and coronal (bottom) planes. (E) Location of an example cell (blue) in each of two flies (top and bottom, respectively) negatively correlated with running. (F) Corresponding activity traces for cells indicated in ‘E’ for each fly. Ethograms shown below for reference. (G) Distribution of behavior time constants (*τ*) vs model *r*^2^ for all flies (N = 18, cells significantly active during behavior in blue, all other cells in gray). (H) Distribution of *τ* vs distribution of time shifts (*ϕ*) for all flies (N = 18), all cells significantly active during behavior. (I) Downsampled composite spatial map of *τ* for all flies, with large values in yellow and small values in red, viewed in the sagittal (left), transverse (right), and coronal (bottom) planes. Scale bar for all maps is 50 *μ*m.

Cells exhibited a remarkably broad range of preferred filter time constants (Fig. 2A, G). 38% of cells had small time constants (*τ* < 4 seconds), reflecting the similarity of the dynamics of behavior and mean neural activity (Fig. 1E). However, 37% of all cells have *τ* greater than 20 seconds, and the overall distribution is bimodal (Fig. 2G). Thus, the neural relationship to behavior has two timescales that approximate the timescales of running itself (Fig 2G and 1E). The median *r*^2^ does not decrease as *τ* increases, indicating that behavior explains a similar fraction of neural activity in cells with small and large behavioral time constants (Fig. 2G). The temporal shifts in the filters were almost always positive and similar to the filter time constants, such that cells with large time constants also had large shifts (Fig. 2H). The locations in the brain of cells with a given behavior time constant exhibit spatial organization (Figure 2I): some brain regions exhibit predominantly small *τ* and other regions exhibit large *τ*. Neurons with large *τ* cluster in the PI region and in lateral areas on the posterior and anterior surfaces (Fig. 2I). Neurons with small *τ* are distributed throughout the brain but most concentrated near the midline on the dorsoposterior surface (Fig. 2I). This region is primarily composed of neurons innervating the protocerebral bridge and fan-shaped body of the central complex and descending neurons innervating the ventral nerve cord (“CX, DN”, Fig. 2I, Fig. S2). This makes sense, as neurons in these brain regions are involved in orienting and locomotion (32, 33).

### Brainwide neural activity correlates with vigorous but not subdued behaviors

Do all behaviors engage the entire dorsal brain, or is running unique? Grooming and running are both precise directed behaviors but differ in the number of limbs they engage, whereas flailing and running both engage all limbs. We define behaviors engaging all limbs as ‘vigorous’ and behaviors engaging fewer limbs as ‘subdued’. Most neurons are noticeably less active during grooming than running (Fig. 3A). Only 3.0% and 2.1% of all cells have *τ* < 4s and a regression weight greater than 0.02 for front or back grooming, respectively (Fig. 3B-C). Only eight cells across all flies were highly correlated with front grooming (*CC* > 0.5, *τ* < 4s), and only two flies had multiple such cells (Fig. 3D). In both flies, these cells were near the periphery of the imaged volume, potentially accounting for their absence in other flies. To quantify brainwide influence of each behavior, we normalize the variance explained by each behavior by the total time each fly exhibited that behavior, relative to running. Front and back grooming respectively account for only 18% and 9% as much variance in neural activity as running in cells with *τ* < 4s (Fig. 3E, Fig. S3). Our observation that the dorsal brain is not broadly engaged during grooming is qualitatively in agreement with prior work proposing that small ensembles of cells are responsible for grooming (34, 35).

**Figure 3.**
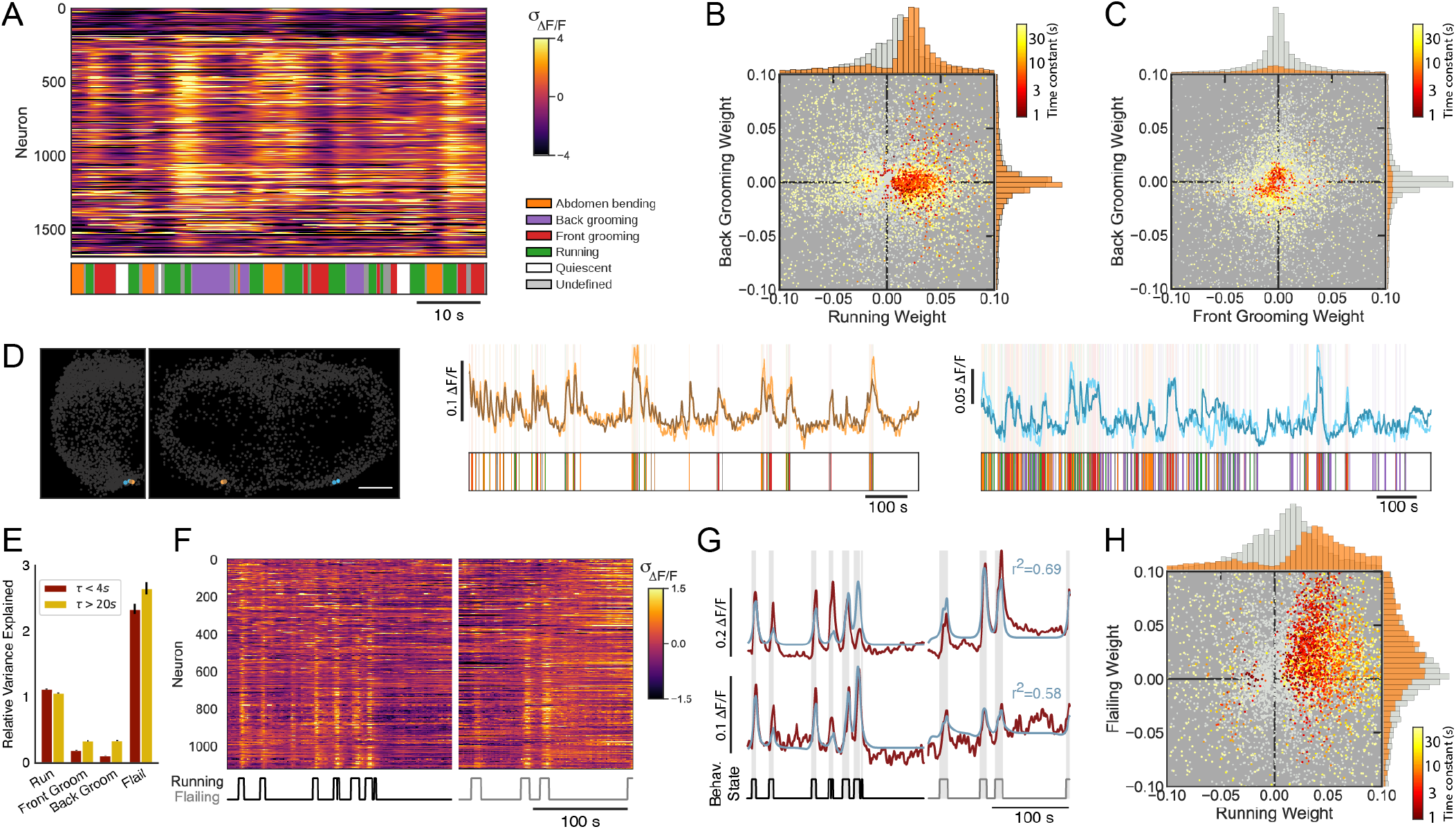
Brainwide neural activity correlates with vigorous but not subdued behaviors. (A) Example raster of z-scored Δ*F*/*F* for all cells from one fly in a short time window, showing individual bouts of many behaviors. Cells ordered by ascending *ϕ*. (B) Regression weights for running vs. back grooming, for all flies (N = 16), all cells significantly active during at least one behavior colored by behavior time constant. Cells not significantly modulated during either behavior shown in gray. (C) Regression weights for front grooming vs. back grooming, for all flies (N = 16). (D) Left, location of pairs of cells in two flies (gold and cyan, respectively) correlated with front grooming. Scale bar is 50 *μ*m. Right, corresponding activity traces for cells indicated at left for each fly. Ethograms shown below for reference. (E) Relative variance explained for each behavior, normalized to running, for all cells and all flies (N = 16 for running and grooming, N = 10 for flailing). (F) Raster of z-scored Δ*F*/*F* for all neurons from a fly running on a spherical treadmill (left) and then flailing in the absence of a spherical treadmill (right), with the timeseries of bouts of activity (running/flailing) shown below. (G) Activity from two example neurons (red) from the same fly as ‘F’, with regression model fits overlaid in blue and behavioral state (running/flailing/quiescent state) shown below. (H) Distribution of regression weights for running and flailing for all cells and all flies (N = 10).

We elicit flailing by removing the treadmill from beneath the fly. The representation of flailing is brainwide and qualitatively similar to that of running (Fig. 3F). 59% of neurons with regression weights greater than 0.02 and *τ* less than 4s during running had equally large regression weights during flailing (Fig. 3G-H). This suggests that global activity does not encode the precise modality of locomotion but rather may encode locomotive vigor or arousal more generally. This is further supported by the observation that unlike grooming, flailing accounts for more variance than running (218% and 262% for *τ* < 4s and *τ* > 20s, respectively. Fig. 3E). Collectively, our results suggest that vigorous behaviors activate global representations, whereas less vigorous or subdued behaviors do not.

### Residual neural activity reveals ensembles of neurons with correlated activity

We next examined the nature of the neural activity not accounted for by our regression model, and thus not grossly linked to any of the identified behaviors. After large-scale locomotion- and other behavior-related activity has been regressed out, the residual activity exhibits rich dynamics across both space and time (Fig. 4A). On average, the fraction of variance explained by behavior (mean *r*^2^ = 0.39) is similar in magnitude to that of the residual dynamics (1−*r*^2^). These residual dynamics include neurons that are highly active during running (Fig. 4B, red, Fig. S4), implying that global and residual activity coexist in the same population of neurons.

**Figure 4.**
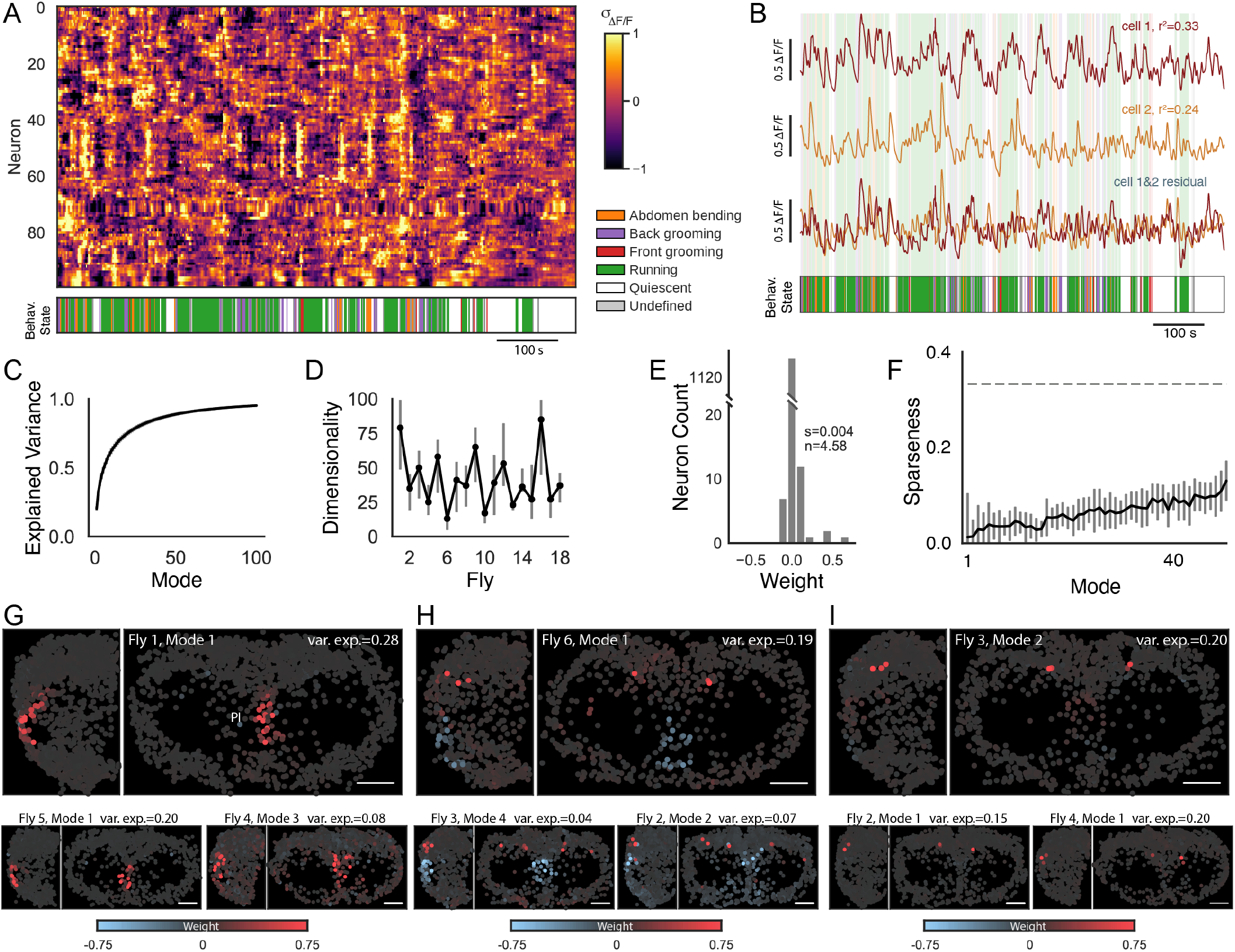
Neural activity not accounted for by behavior is high-dimensional. (A) Example residual of the behavioral regression model reveals rich dynamics and groups of neurons with similar activity (z-scored Δ*F*/*F*, N = 100 cells, ordered by iteratively selecting the neuron most correlated with the previous neuron). Behavior ethogram shown below. (B) Example traces from two selected cells (red, gold, respectively) either before (top, middle) or after (bottom) subtracting the behavioral regression fit, with ethogram shown below. (C) The fraction of total variance explained in the regression residual as a function of the number of PCA modes (mean ± SEM, N = 18). (D) Dimensionality of the regression residual for all flies, calculated as the peak in log-likelihood. Bars indicate ± 1%. (E) Weights of all cells in a single representative PCA mode (fly 3, mode 2). Sparseness = 0.004, corresponding to 4.58 participating neurons. (F) Sparseness of each PCA mode, averaged across all flies (Methods, median ± SEM, N = 18). Dashed line represents Gaussian zero-mean patterns. (G-I) Example maps of weights from leading PCA modes are sparse, approximately symmetric, and exhibit common patterns across flies (scale bar = 50 *μ*m). Shown are examples dominated by Pars Intercerebralis (PI) neurons (‘G’), dorso-posterior neurons (‘I’), and anticorrelations between neurons from the two regions (‘H’). Upper-right in ‘I’ (Fly 3, Mode 2) is the same mode as shown in ‘E’.

We examined the structure of residual activity by performing a principal component analysis (PCA). On average, the first 10 dimensions explain 62% of the residual variance, and subsequent modes each account for no more than 2% of the variance (Fig. 4C). Surprisingly, despite accounting for little variance, many dimensions of activity can be distinguished from noise (41.5 ± 4.6, Fig. 4D, Methods). These PCA modes are very sparse, in some cases involving as few as 4 neurons (Fig. 4E). The average sparseness of the first two modes is 1.3%, meaning that a typical mode involves 18 neurons (Fig. 4F). Thus, each mode explains a small fraction of the total variance but describes a reliable pattern present in neural activity. Counterintuitively, dominant modes are sparser than less dominant modes (Fig. 4F). This suggests that the most reliable patterns in the data tend to contain fewer neurons.

Each PCA mode is sparse and therefore dominated by the activity of a small group of neurons. These modes show spatial organization; for example, small groups of bilaterally symmetric neurons dominate the largest PCA modes (Fig. 4G-I). These modes are similar across flies, although there is variability in which mode explains the most variance in a given fly (Fig. 4G-I). The dominant modes identified by this analysis correspond to ensembles of approximately 20 cells that may comprise functional units. The ensembles often display symmetry across hemispheres. Each functional group is likely to be made up of multiple clusters with even smaller numbers of neurons, perhaps corresponding to specific cell types.

### Residual activity is similar in running and quiescent states

What is the relationship between global behavior-related activity and the sparser residual patterns of activity? One possibility is that residual dynamics could depend on behavioral state so that, for example, a particular residual dynamic pattern only appears during running (Fig. 5A, model 1). Alternatively, residual dynamics could be present in multiple behavioral states but in different state-dependent forms (Fig. 5A, model 2). Finally, residual activity could be independent of behavioral state, and therefore similar, for example, in the running and the quiescent states (Fig. 5A, model 3). We find that the third of these possibilities most accurately accounts for our data; residual activity shows no obvious relationship to behavioral state (Fig. 4A).

**Figure 5.**
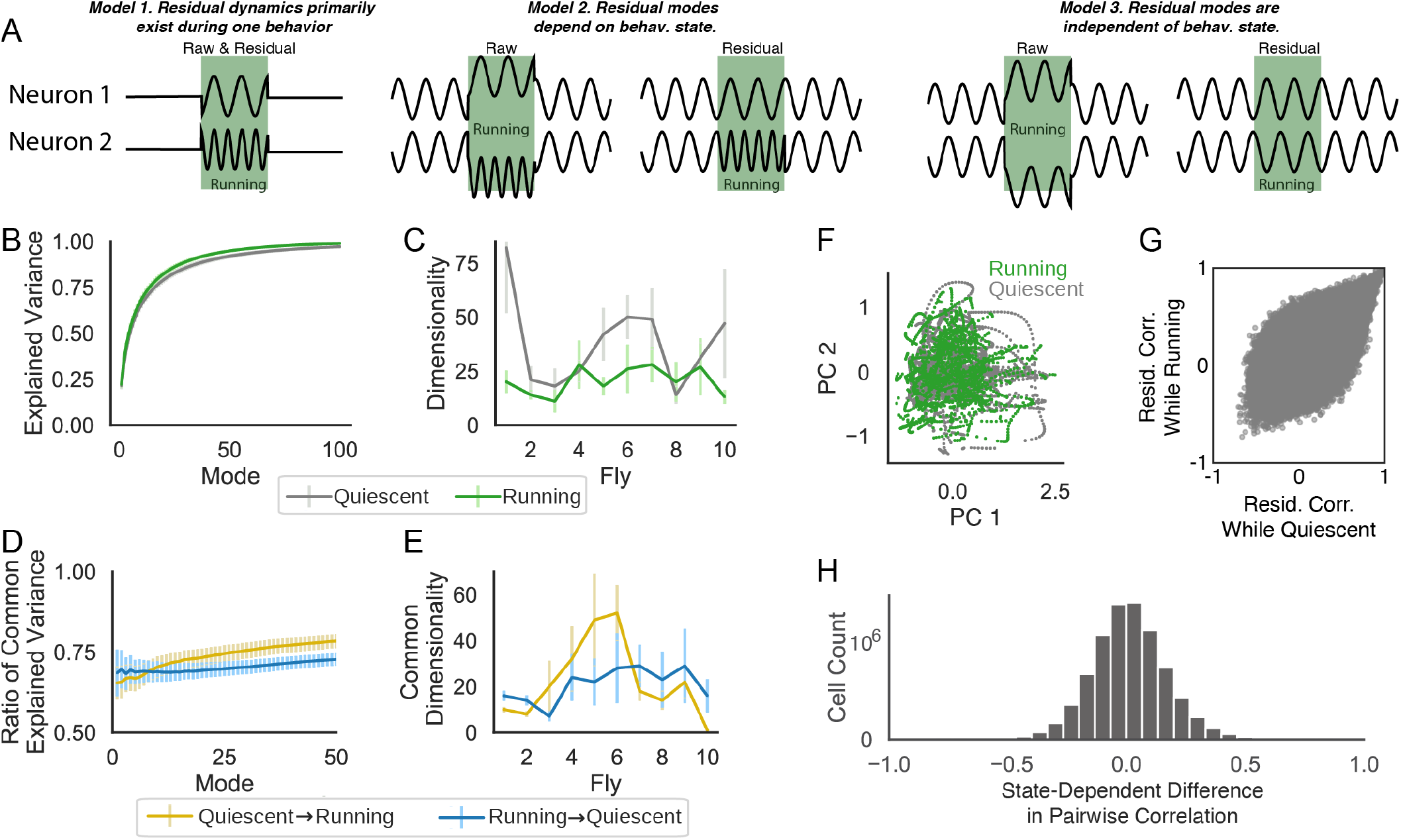
Residual neural activity is largely independent of behavioral state. (A) Possible relationships between residual activity and behavioral state for two cartoon neurons. Model 1: Residual dynamics only exist during one behavioral state. Model 2: Both raw and residual dynamics depend on behavioral state. Model 3: Residual dynamics are independent of behavioral state. (B) Fraction of total variance explained, as in Fig. 4C, but fitting and testing exclusively on times the fly was quiescent or running (gray and green, respectively). (C) Estimated dimensionality for the quiescent and running states, calculated as in Fig. 4D. (D) Using PCA modes calculated as in ‘B’ but evaluating them on the opposite behavioral state. Cumulative variance explained in the opposite behavioral state is divided by variance explained in the fitted behavioral state. (E) Shared dimensionality of the quiescent and running states, calculated as in Fig. 4D. (F) Projection of residual dynamics during the running (green) or quiescent (gray) states onto the first two PCs of the running state for an example fly. (G) Residual pairwise correlation during either the quiescent or running state, for all cells from one fly. (H) Distribution of differences of residual pairwise correlations between the quiescent and running states for all flies (N = 10).

We examined the residual neural activity during a behavioral state (a “subspace”) and compared the subspaces of the running and quiescent states. The amount of variance explained by each mode appeared virtually identical in the two states (Fig. 5B). The dimensionality of these two subspaces is qualitatively similar, but on average the quiescent state is higher dimensional (37.9 ± 6.1) than the running state (20.5 ± 2.0, Fig. 5C). This implies that the running and quiescent states are both complex.

We next asked if the residual activity during the running and quiescent states are not only similar in their complexity but also contain similar dynamics. We therefore determined whether the PCA modes defined in one state explain appreciable variance in the other state. PCA modes defined by activity during the quiescent state explain approximately 75% as much variance in the running state, and PCA modes of the running state explain 75% of the quiescent state (Fig. 5D). This implies that the subspaces occupied by the dynamics in each state are highly overlapping. Furthermore, the di-mensionality of this overlap is similar to the dimensionality of the activity (Quiescent-to-Running = 22.6 ± 5.1, Running-to-Quiescent = 20.8 ± 2.2, Fig. 5E). Moreover, projections of the residual dynamics from both states onto the first two modes of the running state are highly intermingled (Fig. 5F, also see Fig. S5). Collectively, these results indicate that the temporal and spatial structure of the residual activity is similar in the running and quiescent states.

PCA identifies patterns in the correlations across the full population of neurons. To look for state-dependent effects in small groups of cells, we compared correlations between the residual activity of all pairs of cells in the quiescent and running states. These correlations are similar with no large outliers (Fig. 5G-H). Thus, behavioral state and the global pattern of activity associated with it appears to have only a modest effect on the structure of residual activity. This is true not only for the residual dynamics of large populations of neurons but also for the residual correlations between all pairs of neurons (Fig. 5G-H). Thus, behavioral state and residual dynamics appear remarkably independent (Fig. 5A, model 3).

### Cluster analysis reveals spatially segregated groups of neurons with correlated activity

PCA revealed ensembles of spatially organized and functionally related neurons in the residual activity. We identified smaller clusters of correlated neurons by performing hierarchical clustering analysis on the residual activity (Fig. 6A). This procedure builds a tree of similarity between the activity patterns of all cells, where at each branch point the ‘children’ describe potentially meaningful subsets of a given ‘parent’. To look for structure in the data at all spatial scales without defining arbitrary parameters for the number of expected clusters, we identified significant clusters using cross-validation (Methods). Specifically, we determined whether the variance of each child cluster was significantly smaller than the variance of random samples of the same size extracted from the parent cluster (Methods). In this way, we determined whether a given small group of neurons defined a cluster unique from other members of the parent cluster. Both a child and its parent cluster can be significant, and therefore clusters may participate in groupings on multiple scales.

**Figure 6.**
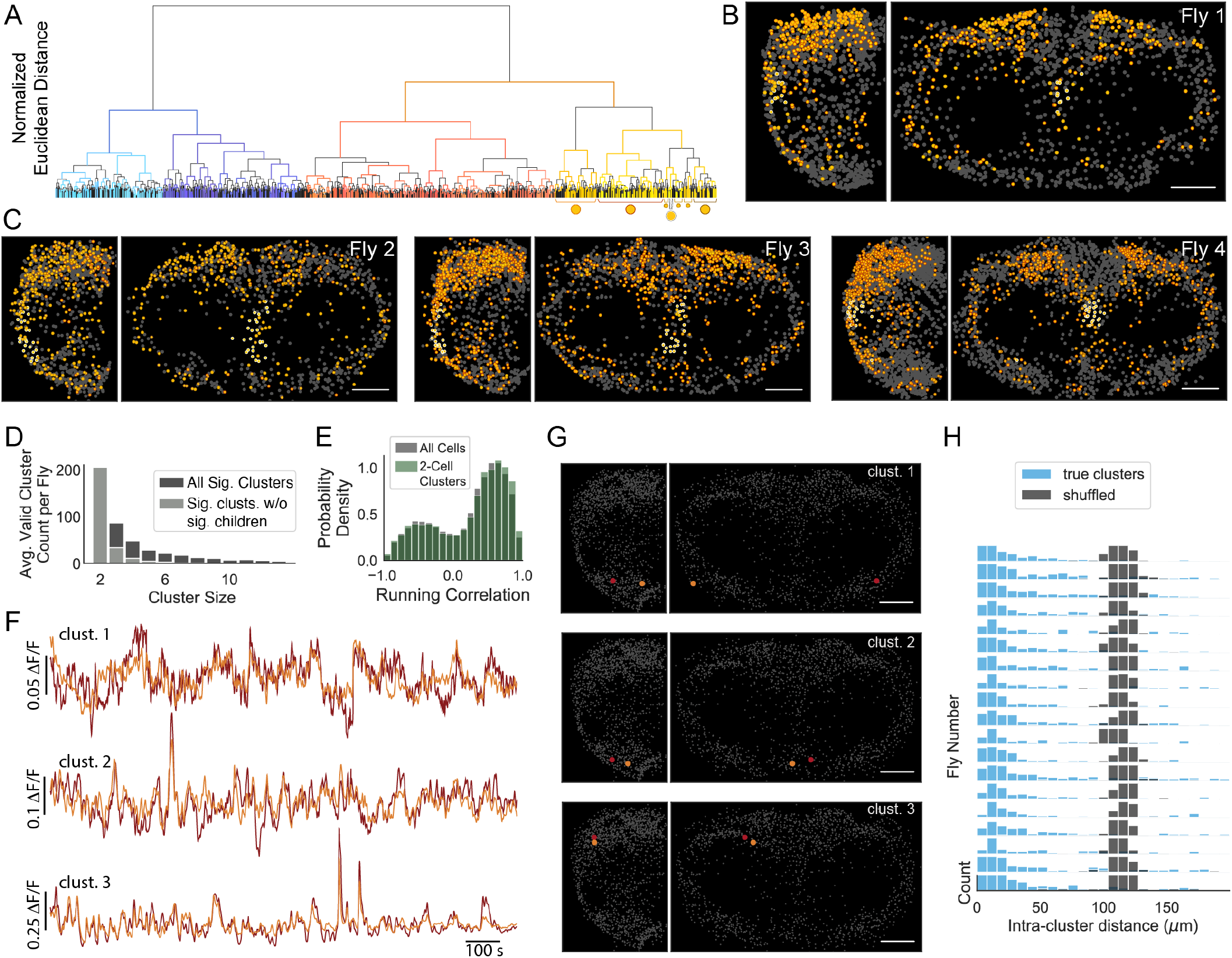
Residual neural activity is composed of organized clusters on multiple spatial scales. (A) The relationship between all cells in one fly, defined by hierarchical clustering on residual neural activity. Vertical axis reflects relative Euclidean distance in activity space. Significance of each cluster was assessed by comparing the variance of the child cluster to the variance of samples from the parent cluster on held-out time points. Branches from non-significant clusters colored black, branches from significant clusters in other colors. Markers under the tree indicate clusters highlighted in ‘B’. (B) Example map of cluster identity for PI cluster and neighboring clusters, with identity indicated by markers in ‘A’. (C) Same as ‘B’ for three additional flies. (D) Distribution of the size of all significant clusters (dark gray) and significant clusters that have no significant children (light gray). (E) Distribution of correlations with running for all cells (gray) and cells belonging to a significant two-cell cluster (green). (F) Residual neural activity from three example clusters each comprising two neurons. (G) Cells belonging to clusters shown in ‘F’ in red and gold, with non-member cells in gray. Scale bar is 50 *μ*m. (H) Euclidean distance between cells belonging to a 2-member cluster (blue), versus randomly assigned cluster labels (gray). Distance computed after superimposing the left and right hemisphere by folding at the midline. Each row shows a different fly and bar height is capped at 30.

Figure 6A shows the full clustering tree for one fly, with each branch colored according to whether the parent was a significant cluster (not significant in black, all other colors significant). We first asked whether significant clusters are spatially organized. A subset of Pars Intercerebralis neurons located near the midline form a spatially compact cluster that is identifiable across flies (Fig. 6B-C, yellow/white). Significant clusters that share a parent with the Pars Intercerebralis cluster are predominantly in posterolateral regions (Fig. 6B-C, orange). Thus, there is spatial organization and stereotypy at multiple spatial scales. The full distribution of sizes for all significant clusters (Fig. 6D) reveals a large number of significant clusters with 2 members. These clusters exhibit diverse residual dynamics, but each cluster consists of pairs of cells with similar dynamics (Fig. 6F). Despite these clusters being defined by residual dynamics, neurons in the same cluster have a similar relationship to global activity and behavior (Fig. S6). As a population, cells within a cluster exhibit a distribution of behavioral time constants and correlations indistinguishable from the distributions across all cells (Fig. 6E, S6). Thus, clusters are highly diverse and participate in the global behavioral state.

By visual inspection, many small clusters appear to be either bilaterally symmetric or spatially localized (Fig. 6G). To quantify this observation, we analyzed the spatial organization by calculating the distance between cells in a cluster after superimposing the left and right hemisphere by folding at the midline (Methods, Fig. S6). Most clusters with two members were more spatially organized than expected by chance (Fig. 6H, S6). The presence of small clusters that are predictive of both activity patterns and spatial location is consistent with the association of cluster identity with function - cells with similar dynamics and similar function are likely to be in similar locations. These observations suggest that the fly brain is composed of many small subpopulations that collectively account for the high dimensionality of the brainwide data. Two-member clusters are embedded in larger ensembles of neurons, implying that the functional relationship between neurons is hierarchical. This is consistent with known classes of cells in the fly brain - for example, Kenyon cells can be subdivided into *α*/*β, α*′/*β*′, and *γ* subclasses, and peptidergic neurons contain subclasses such as insulin-expressing neurons.

Our functional profiling of the brain offers a novel and complementary method of identifying cell types throughout the brain. The vast majority of cells in the central brain can be transcriptionally characterized as consisting of a few thousand distinct cell types that come in clusters of 1-10 neurons per hemibrain (36). Histograms of the number of cells within each cell type from genetic and connectomic cell-typing (36) show an exponential shape similar to that revealed by our activity-based analysis (Fig. 6D). Thus, the smallest spatially organized subpopulations we identified functionally may correspond to genetically defined cell types.

## Discussion

We used SCAPE microscopy to record from a substantial volume of the dorsal brain with cellular resolution, complementing large-scale studies of neuropil regions in the fly brain (6–8, 17, 37). SCAPE imaging permitted us to record from all neurons in a contiguous and large brain volume during spontaneous behavior and revealed global activity that correlates with behavior. SCAPE permits high speed volumetric imaging with cellular resolution, providing an extensive picture of neural activity in space and time. When placed on a ball, flies run, groom, or are quiescent. When suspended, flies often flail. Running and flailing engage a large fraction of the neurons in the imaged volume. A much smaller fraction of the neurons exhibit activity correlated with grooming. A regression model reveals neural activity correlated with running on both short and long time scales. This suggests that most neurons are correlated with the act of running, and a significant fraction are correlated with the tendency to run. Moreover, cells with a given behavioral time constant are spatially organized, in some cases aligning with areas known to be involved in metabolism or locomotion.

Subtracting the dominant activity correlated with behavior reveals additional rich dynamics across time and space. Interestingly, this activity shows little dependence on locomotive state: residual activity exhibits similar spatiotemporal patterns in running and quiescent states. This activity is high dimensional and sparse. Hierarchical clustering reveals small groups of neurons with highly correlated activity, at the extreme comprised of only 2 cells. These functionally defined clusters may correspond to genetically defined cell types in the fly brain. These small circuits do not operate in isolation. Clusters defined by the residual activity also participate in the global behavior-related dynamics. Thus, global patterns may inform local computation and in turn, local computations may influence global patterns.

The global scale of neural activity correlated with locomotion in flies is consistent with findings in worms (2, 3, 14), zebrafish (4, 5, 16) and mice (11–13, 15). Studies in flies (6–9, 17) and those in other organisms pose the question of the mechanism and function of broadly distributed brain-wide activity. In the fly, small identified circuits that control specific behaviors have been elucidated. However, we have shown that most neurons in the fly brain are active during running and flailing, either as actors or observers. This suggests that neurons engaged in specific behaviors, such as mating, aggression or even egg laying, are also active during spontaneous running, without the act of running triggering these other behaviors. Conversely, local circuits controlling behavior may broadcast information globally, generating the activity we observe. This global broadcast could arise from widespread neuromodulation. Alternatively, the recurrent connectivity of the fly nervous system could provide a pathway for this global activity. One such example is the extensive afferent input to the brain from the ventral nerve cord – approximately 2,500 neurons originating in the ventral nerve cord project diffusely to the central brain (33).

We observe a small but substantial fraction of neurons that correlate with locomotion on timescales longer than the duration of individual running bouts. These neurons may represent a locomotor state, the tendency to run. Many of these neurons reside in large posterolateral clusters and in the dorsomedial Pars Intercerebralis. The PI is a predominantly peptidergic domain, and neurons in this region are poised to have influence over extended durations (31). Recent work has implicated a relationship between brainwide behavior-related activity and metabolism (8). Our observation that neurons involved in regulating metabolism are also modulated by running, albeit in a manner distinct from most other neurons, suggests that the causality of this relationship may be bidirectional.

Why does locomotor behavior have privileged access to virtually all neurons in the fly brain? Neurons in multiple neural pathways would likely benefit from knowledge of current behavior. This activity may modulate ongoing behavior or recapitulate past or even predict future behavioral action. In the visual system, for example, locomotion enhances gain and elicits activity in area V1 of mice (22, 25) and in the optic lobe of flies (20, 21, 24). Locomotive behavior in the fly signals not only to primary sensory areas, but to deeper sensory structures such as the mushroom body (38). Moreover, locomotor state in the fly combines with self-generated visual feedback to control posture (39). Efference copies from motor systems to multiple circuits enable the cancellation of self-generated sensory input (40). In artificial intelligence, the utility of proprioceptive feedback to higher-order networks has been demonstrated – in artificial agents trained to solve a variety of tasks, subnetworks charged with representing abstract quantities such as value benefit from knowledge of the agent’s behavior (41, 42). Interestingly, artificial neurons in such subnetworks also tend to have activity correlated with the behavior itself (41). Therefore, locomotor state may provide a useful behavioral context for other computations throughout the brain and it is perhaps not surprising that it elicits the most prominent activity throughout the brain. In short, it is good to know what you are doing.

## Acknowledgements

We would like to thank Tanya Tabachnik for design and manufacturing of our fly treadmill, Barry Dickson for generously sharing fly stocks, Armaan Ahmed, Virginia Devi-Chou, and Benjamin Lucero for assistance processing data, and members of the Axel, Hillman, and Paninski labs for helpful comments and suggestions. This work was supported by grants from the the Simons Foundation (481778, ESS; 542951, LFA, EMCH, RA; 543023, LP), the NSF Graduate Research Fellowship Program DGE 16-44869 (NM), BRAIN Initiative Awards UF1NS108213 and U01NS094296 (EMCH), the NSF NeuroNex Award DBI-1707398 (LFA, LP), the Gatsby Charitable Foundation (LFA), and the Howard Hughes Medical Institute (RA).

## Methods

### Genetics and fly rearing

We imaged female 4-7 day-old flies of the following genotype: w/+; UAS-nls-GCaMP6s/+; Nsyb-Gal4/UAS-nls-DsRed. UAS-nls-GCaMP6s was a gift from Barry Dickson.

### SCAPE light-sheet imaging

Imaging was performed on a SCAPE 2.0 system (27). In brief, the laser sheet was directed through an upright mounted 20x/1.0NA water immersion objective. Emitted light from the sample was separated into two channels by an image splitter outfitted with two dichroic filters and the detected red and green channels were recorded side-by-side on the camera chip. The imaging speed for these experiments was between 8-12 volumes per second, typically covering a volume of approximately 450 × 340 × 150 *μ*m^3^.

### Imaging and behavior preparation

We mounted flies to a customized holder consisting of a 3D printed holder and a laser-cut stainless-steel headplate. We use a spherical treadmill similar to prior designs (43). We monitor the behavior of the fly at 70 Hz, illuminated by 750 nm LEDs using a Basler acA780 camera outfitted with a VZM-450i lens (Edmund Optics) and a near-IR longpass filter (Midwest Optical LP780-22.5, Graftek Imaging). Depictions of the preparation made in BioRender (44).

### Motion correction

To perform image registration of our volumetric imaging dataset, we used the NoRMCorre algorithm (45) augmented with an annealing procedure in which the grid size and the range of permitted local displacements gradually decrease with each iteration. At each step, we computed displacements using the activity-independent DsRed channel and applied the inferred displacements to the GCaMP channel.

### Source extraction and deconvolution

ROIs are defined using watershed segmentation applied to the red channel of a temporally-averaged volume, resulting in 1,631 +/- 109 ROIs per animal. After motion correction, most cells have negligible residual motion, but in some datasets a small fraction of cells have motion that is too nonlinear to be addressed with NoRMCorre. To quantify residual motion and eliminate non-stationary cells, we compute the squared coefficient of variation, *CV* ^2^ = Var[Δ*F*/*F*]/Mean[*F*]^2^ from the red channel. Most ROIs (>95%) have *CV* ^2^ ≪ 1, while some have *CV* ^2^ ≫ 1 and are discarded. No cells exhibit *CV* ^2^ ≈ 1 (Fig. S1). This refinement of ROIs yields 1,419 ± 78 stable, singlecell ROIs per animal.

Although this procedure typically reduces motion artifacts to less than 1 voxel for most cells, we further minimize the impact of residual motion by defining the activity of each cell as the ratio of green and red, *F* = green/red. We then define baseline ratiometric fluorescence, *F*_0_ as the best-fit exponential using least absolute deviation (LAD) regression applied to the derivative of *F*. LAD regression confers robustness to outliers, and working with the derivative of *F* confers robustness to long-timescale nonstationarity. We find similar but slightly noisier activity using simple Δ*F*/*F*_0_ defined on the green channel alone.

### Anatomical alignment across animals

We create a standardized reference frame by coarsely aligning cell locations across flies. Treating every cell as a point, we align the point sets for each brain to a common reference volume using the Gaussian mixture model method developed here: https://github.com/bing-jian/gmmreg.

### Analysis of behavior

We monitor the movement of the spherical treadmill by measuring the total pixel variance between successive frames from the region containing the ball. This unitless estimate of motion aided behavior segmentation, as described below. In some datasets, the spherical treadmill was removed after 10 minutes of imaging. Here, we measured pixel variance in an ROI around the fly’s legs, which provided a measure of a behavior we called flailing, consisting of bouts of rapid leg movements.

We analyze fly behavior both by directly tracking motion of the treadmill (described above) and by tracking eight points on the body of the fly using Deep Graph Pose (28) (DGP; Fig. 1C). We hand labeled the eight selected points in 50 frames from each of 17 videos using the DeepLabCut (DLC) (46) GUI, for a total of 850 labeled frames. We then trained DGP on these frames, which augments the supervised loss of DLC with a semi-supervised loss that incorporates additional, unlabeled frames; we found that this significantly improved the pose estimation, even after post-hoc smoothing of the DLC markers.

We further segment discrete behaviors from the DGP markers using a semi-supervised sequence model (29). We chose to label five salient behaviors commonly observed across all flies: running, front and back grooming, abdomen bending, and a quiescent state. We labeled up to 1,000 frames for each of the five behaviors for each of 20 flies using the DeepEthogram GUI (47), resulting in a total of 33,756 hand labels (quiescent = 6,250, run = 4,950, front groom = 5,700, back groom = 5,480, abdomen bend = 11,376). We supplemented this small, high-quality set of hand labels with a large, lower-quality set of “weak” labels computed using a simple set of heuristics (see details below).

#### Semi-supervised behavioral segmentation

We train a semi-supervised behavioral segmentation model that classifies the DGP markers into one of the five available behavior classes for each time point. The model’s loss function contains three terms: (1) a standard supervised loss that classifies a sparse set of hand labels; (2) a weakly supervised loss that classifies a set of easy-to-compute heuristic labels; and (3) a self-supervised loss that predicts the evolution of the DGP markers. Let x_*t*_ denote the DGP markers at time *t*, and let **y**_*t*_ denote the one-hot vector encoding the hand labels at time *t* such that the *k*^th^ entry is 1 if behavior *k* is present, else the entry is 0. We assume that the hand labels are only defined on a subset of time points 𝒯 ⊆{1, 2, …*T*}. The cross-entropy loss function then defines the supervised objective (ℒ_super_) to optimize:

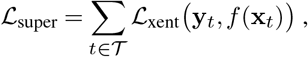

where *f* () denotes the sequence model mapping the DGP markers to behavior labels. We now introduce a set of heuristic labels 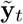, defined at each time point. Computing the crossentropy loss on all time points that do not already have a corresponding hand label defines the heuristic objective:

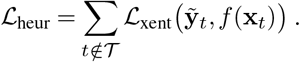

The self-supervised loss requires the sequence model to predict x_*t*+1_ from x_*t*_. To properly do so we now expand the definition of the sequence model *f* () to include two components: an encoder *e*(), which maps the behavioral features x_*t*_ to an intermediate behavioral embedding z_*t*_; and a linear classifier *c*() which maps z_*t*_ to the predicted discrete labels (*ŷ*_*t*_ = *c*(*e*(x_*t*_)). We can now incorporate the self-supervised loss through the use of a predictor function *p*(), which maps z_*t*_ to x_*t*+1_, and match x_*t*+1_ to the true behavioral features x_*t*+1_ through a mean square error loss ℒ_MSE_ computed on all time points:

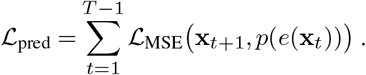

Finally, we combine all terms into the full semi-supervised loss function:

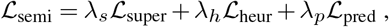

where the *λ* terms are hyperparameters that control the contributions of their respective losses. Note that setting *λ*_*h*_ = *λ*_*p*_ = 0 results in a fully supervised model, while *λ*_*s*_ = *λ*_*h*_ = 0 results in a fully unsupervised model.

For the encoder and predictor networks *e*() and *p*() we use a dilated Temporal Convolutional Network (dTCN) (48), which has shown good performance across a range of sequence modeling tasks. Both networks use a two-layer dTCN with a filter size of 9 time steps and 32 channels for each layer, with leaky ReLU activation functions, and weight dropout with probability p = 0.1. We use 10 fly videos for training and 10 for testing. All models are trained with the Adam optimizer using an initial learning rate of 1*e*-4 and a batch size of 2,000 time points. For the training flies, 80% of frames are used for training, 20% for validation. Training terminates once the loss on validation data begins to rise for 20 consecutive epochs; the epoch with the lowest validation loss is used for testing. To evaluate the models, we compute the F1 score - the geometric mean of precision and accuracy - on the hand labels of the 10 held-out test flies. We average the F1 score over all behaviors and choose the hyperparameters *λ*_*h*_ and *λ*_*p*_ based on the highest score. We then retrain the model with those hyperparameter settings using all 20 flies to arrive at our final segmentation model. We also performed a small hyperparameter search across the number of layers, channels per layer, filter size, and learning rate, and found that our results are robust across different settings (data not shown).

#### Heuristic labels

The addition of a large set of easily computed heuristic labels improves the accuracy of the behavioral segmentation (29). Below, we provide more detail on these heuristics. Note that we choose conservative values for the thresholds in order to decrease the prevalence of false positives. A consequence of this choice is that some time points are not assigned a heuristic label; nevertheless this procedure adds enough high-quality information to substantially improve the models.

##### Run

We first estimate the time points at which a fly is running by utilizing the treadmill motion energy (ME). We transform the treadmill ME to lie in the range [0, 1], then assign the ‘run’ label to time points when the treadmill ME is above a threshold (0.5).

##### Quiescent

We compute the average ME over all DGP markers for each time point, then denoise this one-dimensional signal with a total variation smoother (the `denoise_tv_chambolle` filter from the sklearn (49) Python package). We then transform this signal to approximately lie in the range [0, 1] (the 99th percentile is mapped to 1 in order to make this process robust to outliers). We assign the ‘quiescent’ label to time points when this signal is below a threshold (0.02) and the fly is not running (according to the previous heuristic).

##### Abdomen bend

We compute the average ME over the abdomen markers, then denoise this signal and transform it to approximately lie in the range [0, 1]. We assign the ‘abdomen bend’ label to time points when this signal is above a threshold (0.9) and the fly is not still or running according to the previous heuristics.

##### Front and back groom

We compute the average ME over the forelimb markers, then denoise this signal and transform it to approximately lie in the range [0, 1]. We assign the ‘front groom’ label to time points when this signal is above a threshold (0.05), the corresponding back groom signal (computed from the hindlimb markers) is below a threshold (0.02), and the fly is not still, running, or bending its abdomen according to the previous heuristics. We assign the ‘back groom’ label in an analogous manner.

### Regression model

We regressed each neuron’s activity against all behaviors (*B* = {running, front grooming, back grooming, flailing}) filtered using a fitted time constant (*τ*_*i*_) and temporal shift (*ϕ*_*i*_) unique for each cell (*i*). Thus, we model the activity *f* of cell *i* at time *t* as

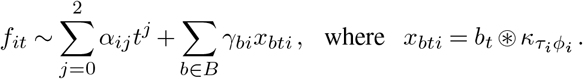

The *γ* coefficients describe the relative importance of each behavior in accounting for the activity of each cell, while the *α* coefficients capture drift independent of behavior. The convolution kernel is 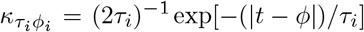. This symmetric kernel is useful for disentangling smoothness from causality. A cell with a broad kernel should have |*ϕ*| ≥ *τ*, with the sign of *ϕ* determining the direction of causality. A lag of |*ϕ*| ≈ *τ* should not be interpreted as a true lag, but rather a reflection of causality with smoothness constraints.

### Dimensionality reduction

We performed principal component analysis (PCA) on the residual activity after subtracting the regression model fit. We quantified the dimensionality of this residual activity as the number of PCA modes that maximize the log likelihood of the lower dimensional subspace on held-out data. We fit the principal components on 80% of all time points and evaluate the log likelihood on the remaining 20%.

To quantify the degree of sparseness of PCA modes without selecting a threshold, we calculate the participation ratio of each mode 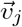 as

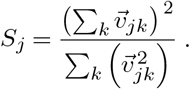

The participation ratio of a zero mean Gaussian vector is approximately 0.33, which is a useful null hypothesis for the existence of either sparse or dense structure in the PCA modes. We sorted residual activity by behavior label and then performed PCA separately on each behavior’s set of time points to quantify the residual subspace (x_*b*_) of each behavior *b*. To compare the subspaces of two behaviors, for example running and the quiescent state, we quantified the common variance explained and the common dimensionality. We defined common variance explained (*E*_*mb*_) for *m* modes as

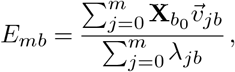

where *b* is the behavior on which the PCA modes were defined, and *b*_0_ is the other behavior. Similarly, we define common dimensionality by cross validating the projection of one subspace onto the modes of the other 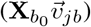.

### Clustering

We performed agglomerative hierarchical clustering on residual neural activity using Euclidean affinity and ward linkage.

To look for structure in the data at all spatial scales without defining arbitrary parameters for the number of expected clusters or an affinity threshold, we identified significant clusters using cross-validation. We performed clustering on 80% of the time points, and evaluated the validity of the identified clusters on the remaining 20%. Specifically, we evaluated the intra-cluster variance on held out time points for each cluster and for size-matched samples from its parent cluster. The number of selected samples was

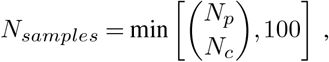

where *N*_*p*_ and *N*_*c*_ are the number of neurons in the parent and child cluster, respectively. A child cluster was deemed significant if its test variance was less than that of the samples (*p <* 0.05). Both a child and its parent cluster can be significant. We defined the intra-cluster distance for each cluster by first folding the brain along the midline thus, the lateral coordinate of each cell was equal to its distance from the midline (Fig. S6). We then compute the Euclidean distance between the coordinates of each cell in a cluster. We performed this analysis on both the identified and randomly shuffled clusters of the same size to validate our results.

## Supplement

**Figure S1.**
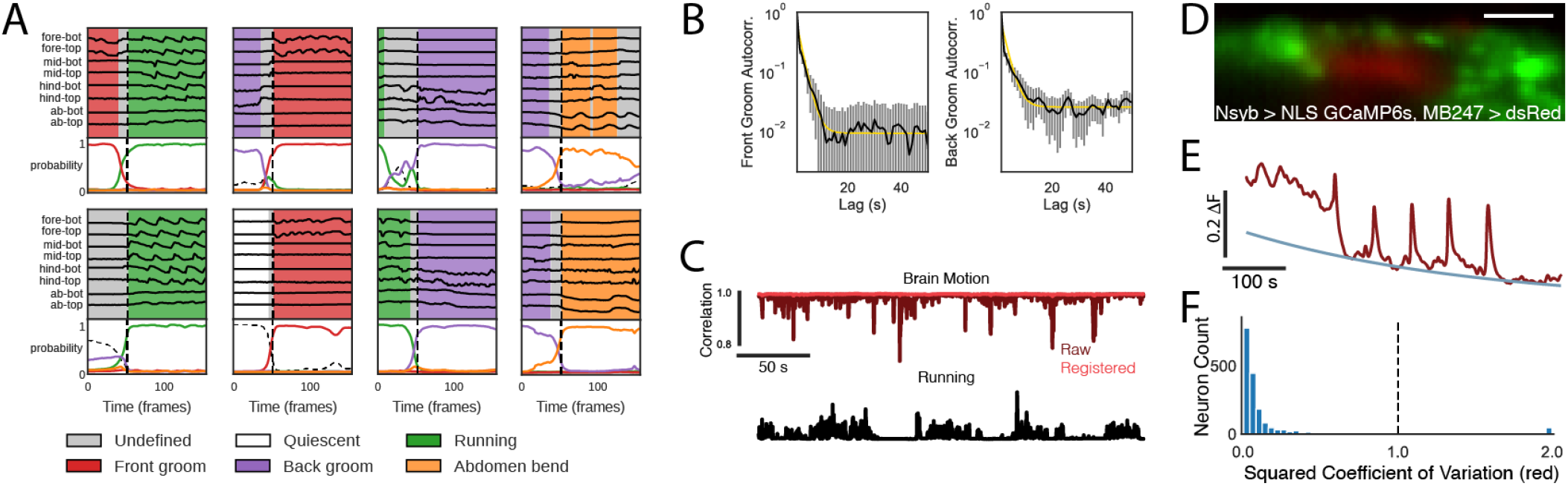
(A) A semi-supervised sequence model (29) extracts a time series of discrete behavioral states from DGP points. Example trajectories of the 8 tracked points shown in black above, ordered from anterior to posterior (fb:front bottom, ft:front top, mb:middle bottom, mt:middle top, hb:hind bottom, ht:hind top, ab:abdomen bottom, at:abdomen top). Inferred probability of each behavioral state is shown below, showing a transition from running to back grooming. (B) The autocorrelation of front grooming (left, black) and back grooming (right, black) are best fit by a single exponential plus a constant offset, with time constants of 2s and 3s, respectively (gold). (C) Motion of the brain volume before (dark red) and after (light red) registration, quantified as the correlation coefficient between red fluorescence and a single template image, compared to running (black). (D) Expression of NLS-GCaMP6s driven by Synaptobrevin (green), with Kenyon Cells labeled in red (MB247>dsRed) for a slice through the calyx in one hemisphere. Scale bar = 20 *μ*m. (E) We define baseline ratiometric fluorescence (blue) as the best-fit exponential using least absolute deviation (LAD) regression applied to the derivative of ratiometric fluorescence. Raw ratiometric fluorescence shown in red. (F) To quantify residual motion and eliminate non-stationary cells, we compute the squared coefficient of variation from the red channel. Cells with values greater than 1 were rare and eliminated.

**Figure S2.**
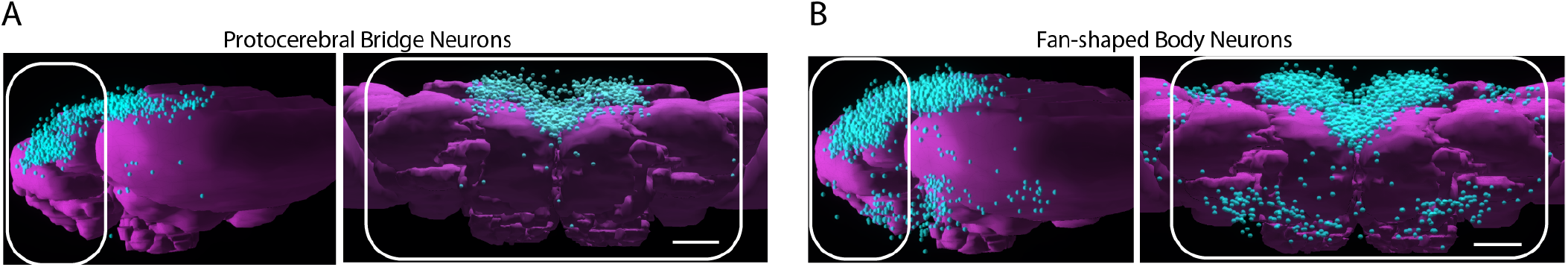
Rendering of cell bodies from all neurons innervating either the Protocerebral Bridge (A) or the Fan-shaped Body (B) of the Central Complex. Renderings created using FlyCircuit (50). In each case, left and right show sagittal and transverse projections, respectively. White rectangles indicate approximate viewing window in our data, and scale bar = 50 *μ*m.

**Figure S3.**
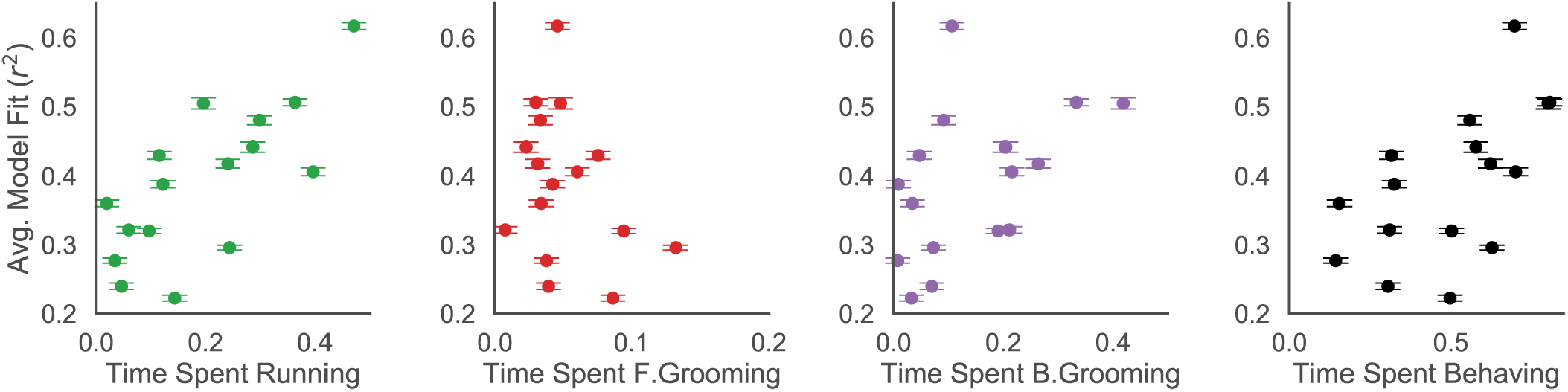
Fraction of variance explained by behavior regression model for each fly versus time spent running (green), front grooming (red), back grooming (purple), or the sum of all behaviors other than the quiescent and undefined states (black). Fraction of time spent running is most predictive of model fit.

**Figure S4.**
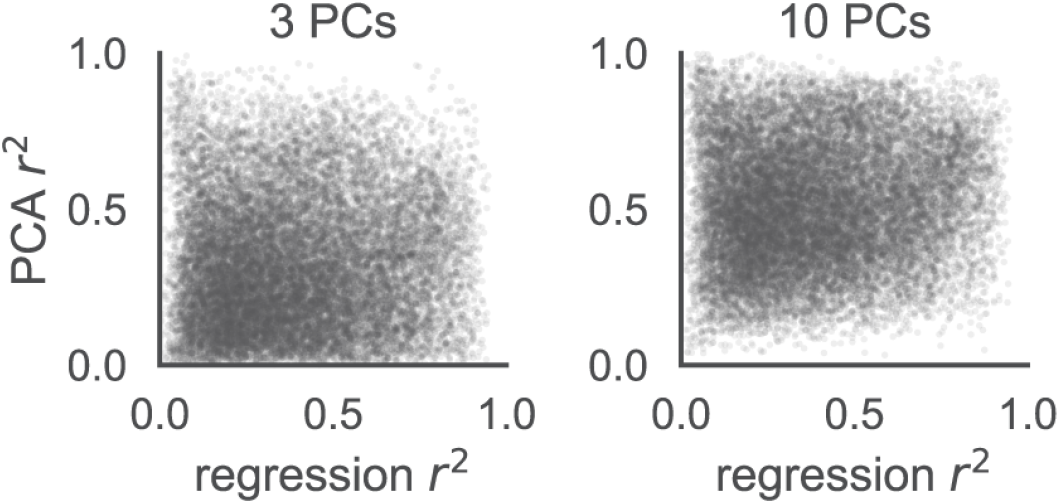
Variance explained by behavior regression model versus residual variance explained by first 3 PCs (left) or first 10 PCs (right) for all cells and all flies (N = 18). Each point is a cell. Variance accounted for by behavior (regression *r*^2^) and variance explained by leading PCs (PCA *r*^2^) are uncorrelated over all cells, implying that global and residual activity coexist in the same population of neurons.

**Figure S5.**
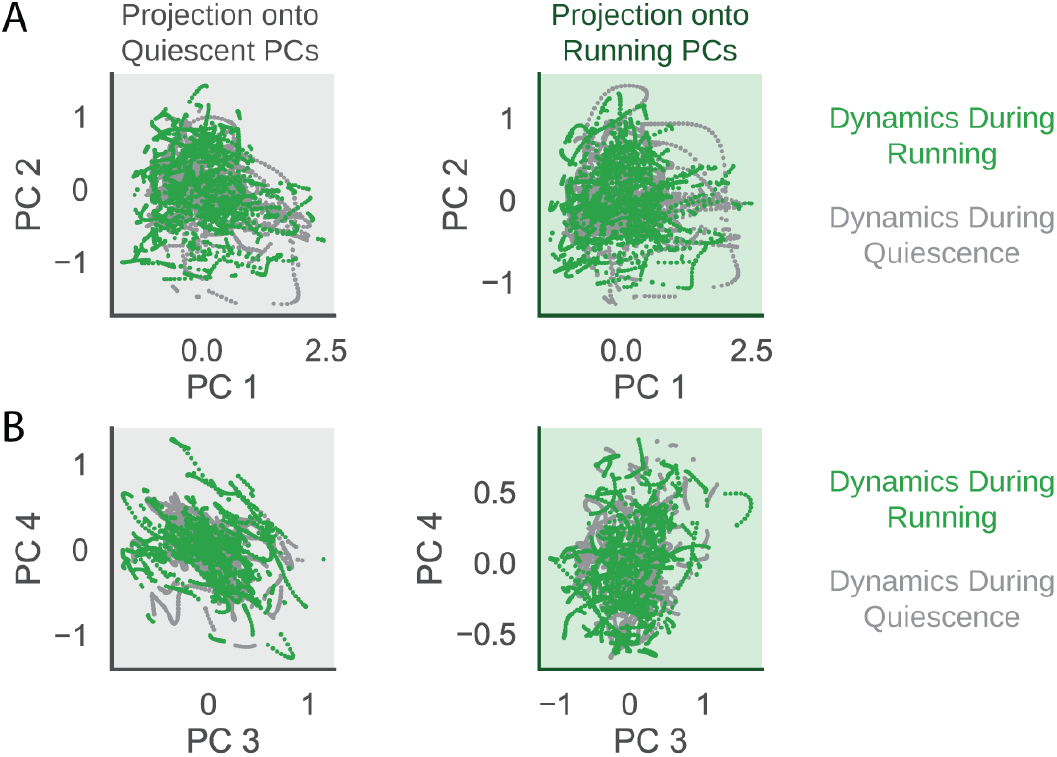
(A) Projections of the residual dynamics during running and quiescence onto the first two modes of the quiescent state (left) and the running state (right) for an example fly. (B) Same as ‘A’, for PCs 3 and 4.

**Figure S6.**
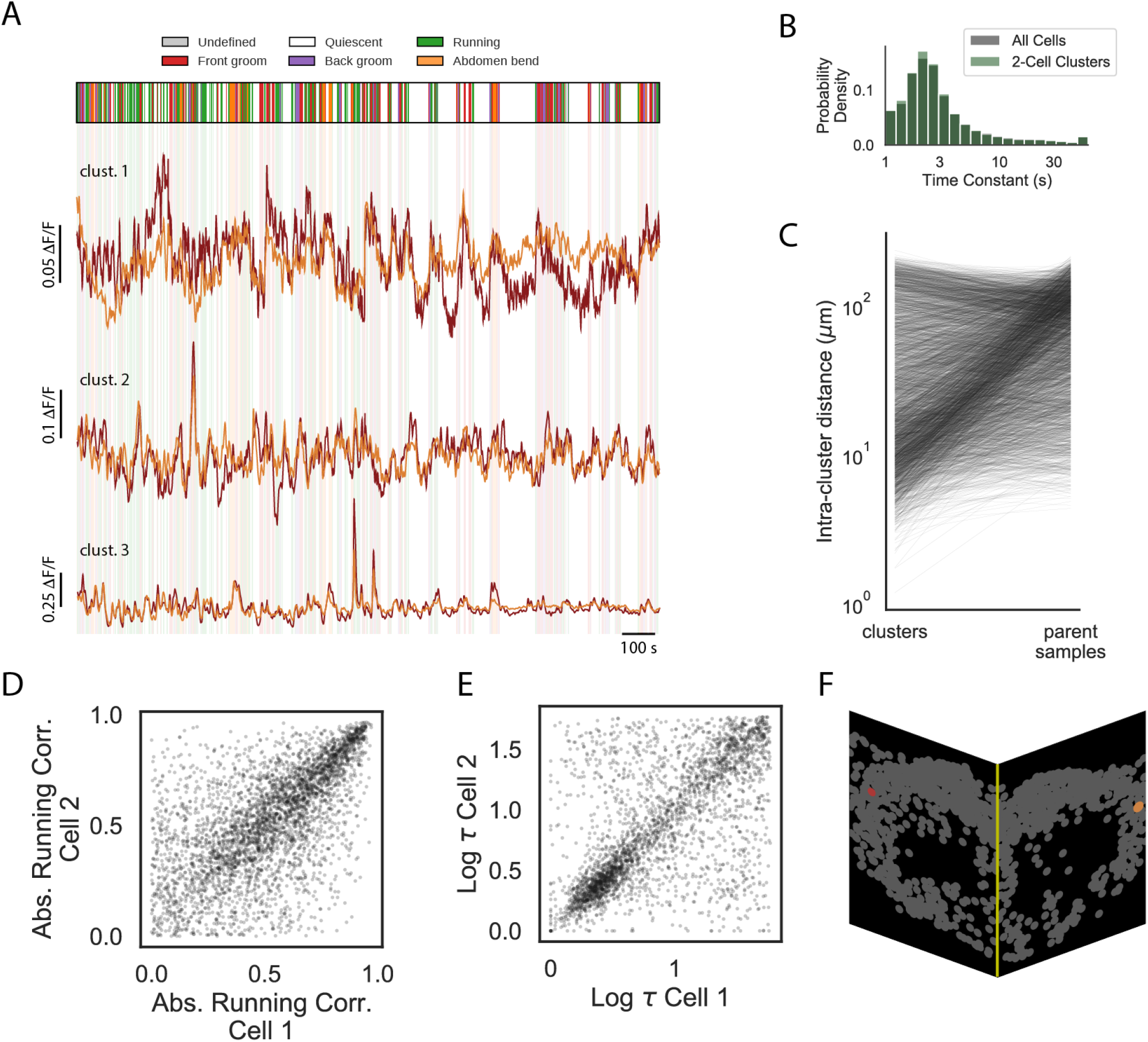
(A) Activity of cells from the three different clusters shown in Figure 6E before subtracting regression fit. (B) Distribution of behavior time constants for all cells (gray) and cells belonging to a significant two-cell cluster (green). (C) Intra-cluster distance of all 2-cell clusters versus the same quantity from samples of each cluster’s parent. (D) Absolute value of correlation with running for each cell in a significant two-cell cluster versus absolute value of correlation with running for its partner cell. (E) Same as ‘D’, for log of behavior time constant. (F) To evaluate spatial organization of clusters, we calculate distance between member neurons after folding the volume along the midline (yellow).

